# Revealing Water-Mediated Activation Mechanisms in the Beta 1-Adrenergic Receptor via OneOPES-Enhanced Free Energy Landscapes

**DOI:** 10.1101/2025.03.03.641221

**Authors:** Simone Aureli, Valerio Rizzi, Nicola Piasentin, Francesco Luigi Gervasio

**Affiliations:** School of Pharmaceutical Sciences, University of Geneva, Rue Michel Servet 1, 1206, Genève, Switzerland; Institute of Pharmaceutical Sciences of Western Switzerland (ISPSO), University of Geneva, 1206, Genève, Switzerland; Swiss Institute of Bioinformatics, University of Geneva, 1206, Genève, Switzerland; Department of Chemistry, University College London, London, WC1E 6BT, United Kingdom

## Abstract

The beta-1 adrenergic receptor (ADRB1) is a prominent pharmacological target due to its critical role in regulating cardiovascular function and is therefore at the forefront of therapeutic interventions in heart diseases. Here we explore the activation mechanism of ADRB1 in both apo (unbound) and holo (adrenaline-bound) forms with OneOPES, a novel multi-replica enhanced sampling simulation algorithm. Our approach leads to converged and reproducible free energy landscapes as shown by independent simulations and identifies key water-mediated interactions that ease structural rearrangements crucial for the activation of ADRB1. The detailed computational analysis provides a comprehensive understanding of the effects of adrenaline on ADRB1’s activation mechanism as well as the role of sodium ions, protonation states and microswitches. Our methodology can be adapted to other ligands and receptors and serves as a blueprint for computational exploration of agonist-induced activation of ADRB1 and other class A GPCRs, paving the way for the development of drugs with fine-tuned modulatory effects.

## Introduction

G-protein-coupled receptors (GPCRs) are one of the largest and most versatile families of membrane proteins, playing an integral role in transducing extracellular signals into cellular responses [1, 2]. They are involved in various physiological processes, including sensory perception, immune response, and neurotransmission [3, 4, 5]. Owing to their central role in numerous signaling pathways, GPCRs have become prominent targets in the pharmaceutical industry, with approximately one-third of all marketed drugs acting on these receptors [6, 7, 8]. In this regard, more than 1000 GPCR structures have been released on the Protein Data Bank (PDB), providing a wealth of structural information [9, 10, 11]. However, experimental high-resolution descriptions of their activation mechanisms are difficult to obtain and have only been reported for a few receptors thanks to NMR [12], time-resolved X-ray crystallography and cryo-EM [13, 14]. Still, the details of GPCRs functional dynamics matter for a full biophysical understanding of their allosteric regulation and importantly for the rational design of more effective and less toxic drugs [8, 15].

To this end, molecular dynamics (MD) simulations have greatly contributed to the understanding of the details of ligand recognition and activation dynamics of GPCRs [16, 17, 18, 19, 20, 21, 22, 23]. Even long MD simulations fall short in reliably capturing the conformational changes associated with receptor activation, due to the limited time-scales that they can sample, underscoring the need for enhanced sampling approaches that can provide a detailed description of the activation mechanisms in their entirety [15, 24].

In this respect, collective variable (CV) based enhanced sampling algorithms, such as Umbrella sampling, Metadynamics [25, 26] and Gaussian accelerated [27] MD have been shown to be particularly effective [28, 24, 29, 30, 31, 32, 19, 33]. However, such methods rely on the definition of effective CVs that capture all the relevant slow degrees of freedom associated with the process of interest. This is far from trivial, making it very difficult to obtain well converged and reproducible free energy landscapes associated with complex functional dynamics such as that involved in GPCR activation [34, 35].

For instance, Metadynamics was used to investigate the binding mode and affinity of adrenaline and noradrenaline toward ADRB1 and ADRB2 [36], revealing distinct binding modes for the two endogenous ligands between these receptor subtypes. However, due to the lack of acceleration of the slow degrees of freedom associated with the receptors’ functional dynamics, the effects of ligand binding on the activation mechanisms of the GPCRs were not explored. To address the difficulty of CV selection for GPCR activation, data-driven CVs have been developed, such as the A100 [24]. PTMetaD [37, 38], a combination of parallel tempering and Metadynamics, has been used to improve convergence problems due to the use of non-optimal CVs [28]. However, even these approaches required very extensive and expensive simulations due to the large number of replicas needed to converge the free energies and not always lead to fully converged free energy landscapes.

We have recently developed a combined enhanced sampling approach called OneOPES to converge free energy landscapes with non-optimal CVs [39]. OneOPES introduces a temperature gradient along a ladder of 8 replicas and accelerates several different CVs simultaneously, allowing accurate and efficient estimation of free energy landscapes with reasonable sampling times. The algorithm was successfully used to converge the free energy associated with the folding of small proteins, molecular recognition, and ligand binding to biochemical targets [40, 41, 42]. So far, however, it has not been used to sample complex conformational changes such as those that are required for the activation of GPCRs.

Here we combine OneOPES with a set of tailored CVs and apply it to study the activation mechanism of one of the most pharmacologically relevant and well-studied members of the GPCR family, the *Beta-1 adrenergic receptor* (ADRB1) [43, 44, 45]. While ADRB1 is primarily involved in the regulation of cardiovascular functions, it is at the forefront of therapeutic interventions for various heart diseases [46, 47]. Drugs that modulate ADRB1 activity, such as beta-blockers, are widely prescribed for the treatment of conditions like hypertension, heart failure, and arrhythmias. Common beta-blockers, including metoprolol, bisoprolol, and atenolol, exert their therapeutic effects by antagonizing ADRB1, thereby reducing heart rate and blood pressure. These drugs are essential in managing cardiovascular diseases and preventing adverse outcomes, such as myocardial infarction and stroke. Given the receptor’s central role in these processes, a deeper understanding of its activation mechanism, particularly in the presence of its endogenous ligands (e.g. adrenaline), is essential for the rational design of more effective drugs.

Combined with a set of CVs specifically tailored to sample the complex events involved in receptor activation, OneOPES demonstrated its ability to converge the free energy landscapes associated with the activation mechanisms of apo ADRB1 and the adrenaline-bound ADRB1 receptor. Independent simulations converged on statistically equivalent free energy landscapes in a reproducible manner and within a reasonable simulation time length. The converged free energy landscapes allowed us to quantify the differences due to the bound agonist, to the presence of a sodium ion in the conserved sodium pocket, and to the protonation state of conserved residue. Moreover, changes in a conserved network of intra-helical water molecules were observed to directly facilitate the structural rearrangements necessary for the receptor’s activation [42], in line with the importance of solvent-mediated allosteric networks emerging from network analysis and GPCR design [48, 49].

Our simulations identified key water-mediated interactions that are integral to the ADRB1’s activation pathway and suggest that water molecules play a crucial role in the structural dynamics of ADRB1, thereby influencing its activation by adrenaline. The insights gained from this study not only enhance our understanding of GPCR activation but also demonstrate the potential of OneOPES in quantifying changes in the population of end-point and intermediate states along the activation pathway due to the binding of agonists. This comprehensive understanding of the activation mechanism of ADRB1 paves the way for the design of novel agonists and antagonists, potentially leading to targeted therapeutic developments in cardiovascular medicine, while the general applicability of our computational approach may lead to the design of better drugs for other GPCR targets.

## Results

### Apo and holo ADRB1 simulations

Our aim is to characterise the structural changes associated with ADRB1 activation and to compute the associated free energy landscape, considering both its apo form and its endogenous ligand (adrenaline) bound form. To this end, we modeled the WT human holo and apo forms from the crystal structures (PDB ID: 7BVQ and 7BTS) [36], and embedded them in a simple lipid bilayer (POPC/CHL 80:20, see Fig. 1a). To enhance the sampling and compute the free energy landscapes, we used OneOPES together with a set of tailored CVs, capturing the many slow degrees of freedom associated with the conformational change (for a detailed description of the CVs and the sampling conditions employed, please refer to the “Methods” section, Tab. S1, and Fig. S1). Each OneOPES simulation of ADRB1 was run independently three times using 8 replicates, with each independent simulation accumulating a total of 16.8 µs of sampling for the apo and 21.6 µs for the holo (see Fig. S2 and S3 for additional details). We also repeated the simulations to quantify the effects of D^2.50^ and the presence of Na^+^ in the conserved sodium pocket.

**Figure 1:**
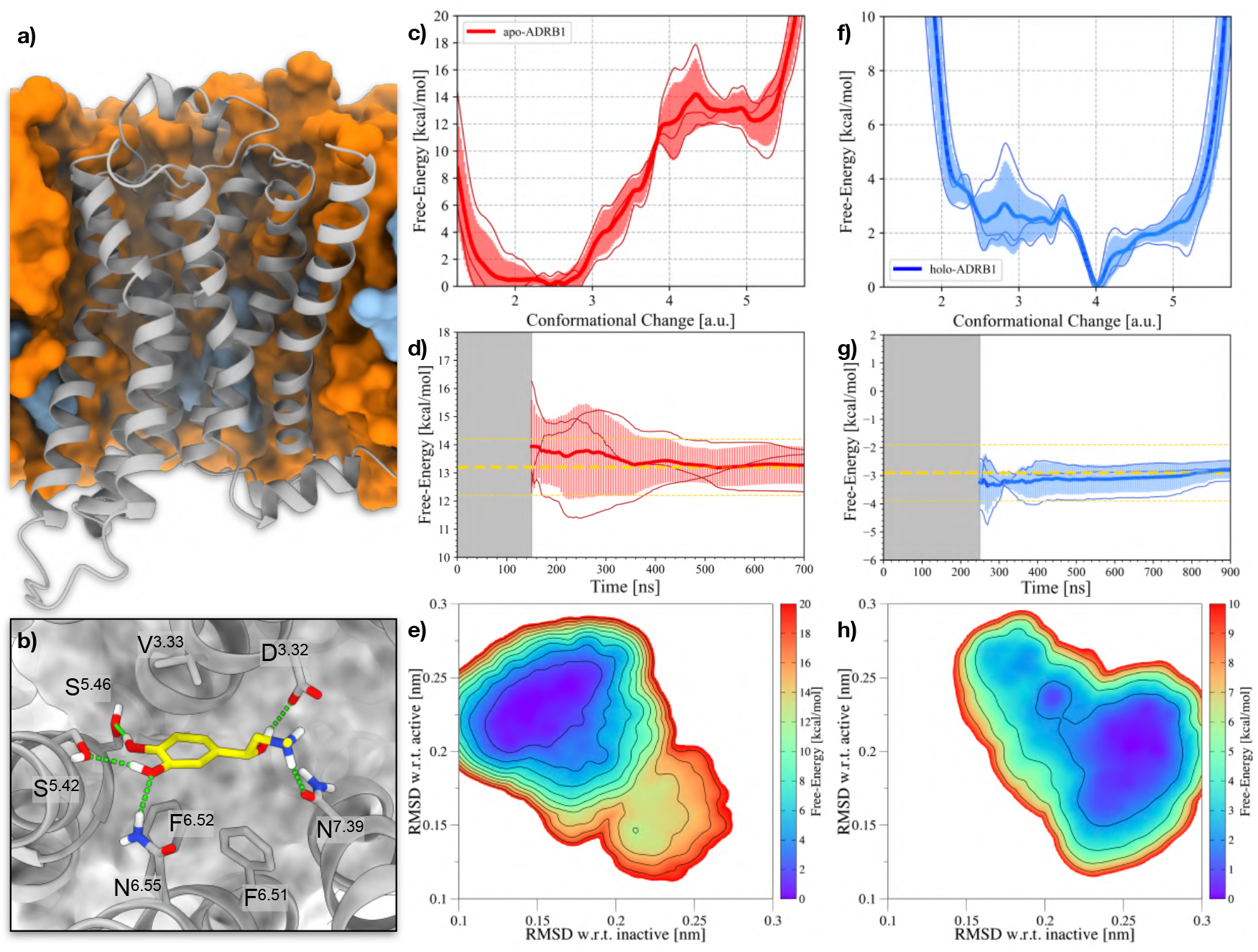
Results of the OneOPES simulations on the apo- and holo-ADRB1 systems. **a)** ADRB1 embedded into a POPC/CHL (80:20) model membrane. ADRB1 is colored in gray, whereas POPC and CHL are depicted in orange and cyan, respectively. **b)** Binding mode of adrenaline (in yellow) bound to ADRB1. **c)** 1D FES as a function of the Conformational Change CV for the apo-ADRB1 systems. **d)** Free-energy difference between inactive and active states in the apo-ADRB1 systems over time. In (c) and (d), the red solid line represents the average of the three replicas, while the transparent red area illustrates the standard deviation. **e)** Average 2D FES concerning the RMSD of both the inactive and active structures for the apo-ADRB1 systems. **f)** 1D FES as a function of the Conformational Change CV for the holo-ADRB1 systems. **g)** Free-energy difference between inactive and active states in the holo-ADRB1 systems over time. In (f) and (g), the blue solid line represents the average of the three replicas, while the transparent blue area illustrates the standard deviation. **h)** Average 2D FES with respect to the RMSD of both the inactive and active structures for the holo-ADRB1 systems. In (d) and (g), the average value is shown as a yellow dashed line. Error bars of 1 kcal/mol appear as yellow dashed lines. In (e) and (h), isolines are drawn every 2 kcal/mol.

To enhance the clarity of the discussion, in the following we present various 1D and 2D projections. For the first and main projection we use an overall “*Conformational Change*” (CC) CV, which is a combination of RMSD from active and inactive structures that is able to track even subtle differences in ADRB1’s plasticity along the functional dynamics (see Fig. S4a and S4b for additional details). Reweighting the accumulated bias potential from the 3 independent simulations onto this CV yields 3 1D FES. In Fig. 1c the curves are shown together with the average and the error bands around the average obtained from the independent simulations. In Fig. 1d the convergence of the ΔG between the active and inactive states is shown. After ∼ 700 ns, all three independent simulations converge to a value of 13 ± 1 kcal/mol. We would like to stress the importance of testing the convergence of the free energy profiles and computing the error bars from independent simulations. This is far from trivial and, so far, due to the significant computational cost, most of the free energy landscapes reported in the literature for GPCR activation, including our own [24, 28], have been obtained from a single run and the error bars from block analysis rather than from multiple independent simulations.

The free energy profile computed for the apo ADRB1 is qualitatively similar to the scheme proposed by Weis and Kobilka [50] and reveals the presence of three distinct regions: a deeper minimum at CC = 2–3, corresponding to the inactive state; a high-energy minimum at CC = 5, representing the active state; and a transition peak at CC = 4, which – based on the status of the micro-switches – we identified as an intermediate state along the activation pathway (see Fig. 1b). The free-energy difference between the inactive and active states is consistently determined from the 3 independent OneOPES simulations to be 13 ± 1 kcal/mol. This value is obtained already after 150 ns of simulation and remains stable until the end of the simulations, providing a quantitative measure of the energetic cost associated with ADRB1 activation (see Fig. 1d).

Our results agree with the mechanism deduced from NMR measurements [12], in particular with respect to the presence of the intermediate (pre-active) state. Our calculated free energy difference for the activation of the apo WT receptor cannot be directly compared with NMR results because, as expected from the high free energy differences, the population of active states is too small to be measured in the experiments. Moreover, the experiments were obtained on a thermostabilized mutant of the turkey ADRB1. In addition, the effect of the reported protonation of the conserved residue D^2.50^, which greatly reduces the free energy difference, must be taken into account (see e.g. Fig.5c).

In the following, the 2D free energy projections presented are computed as an average of the FES from the three independent simulations. Starting from the apo-ADRB1, we reweighted the accumulated bias potential on a 2D space defined by the RMSD with respect to both inactive and active ADRB1 structures (see Fig. 1e). This representation enables the identification of a minimum-energy activation pathway, delineating the re-arrangement ADRB1 must undergo for its activation. We repeated our OneOPES simulations on the holo-ADRB1 system to investigate the impact of a natural agonist, adrenaline, on receptor activation (see Fig. 1b). The 1D FES profile obtained by reprojecting the bias potential onto the CC CV, shows a deep minimum at CC = 4, aligning with the previously identified intermediate state (see Fig. 1f). As expected from NMR measurements, the presence of adrenaline stabilizes this conformational intermediate [12], effectively lowering the energetic cost toward activation, which, over the simulation time, we estimated in a free-energy difference of -3 ± 1 kcal/mol (see Fig. 1g). The ΔG to the fully active state, which as in the apo receptor is found around CC = 5, is less than 2 kcal/mol, in agreement with experiments showing that an increase in the pressure is sufficient to shift the conformational ensemble towards the fully active state [51].

To properly discern the features of the adrenaline-induced intermediate state, we extracted and analyzed the whole conformation pool of structures located at CC = 4. A cluster analysis on adrenaline’s binding mode revealed the presence of a single largely populated cluster, whose position in ADRB1’s orthosteric binding site slightly differs with respect to the crystallographic active state. As shown in Fig. S4c and Fig. S4d, adrenaline is shifted towards TM5 in the intermediate state, a re-arrangement that, while preserving most of the ligand-GPCR interaction network, disfavors the engagement of N^6.55^ with adrenaline’s hydroxy groups. Conversely, N^6.55^ tends to form H-bond with N^7.39^’s side-chain, which also preserves its interaction with adrenaline’s amine. The identification of a novel intermediate structure of ADRB1 where adrenaline preferentially binds may have significant implications for drug discovery and rational design of potent ADRB1 agonists. This intermediate state represents a previously uncharacterized conformation that could serve as a valuable target for structure-based drug design approaches, such as molecular docking and virtual screening. To encourage further exploration and aid drug design efforts, we provide the atomic coordinates of this ADRB1 intermediate structure as part of the Supplementary Material.

The 2D FES, built again using RMSD values relative to the inactive and active ADRB1 structures, shows a significant shift of the activation pathway toward regions of the free-energy landscape that were previously less explored in apo-ADRB1 (see Fig. 1h). Thus, our results provide evidence that adrenaline changes the activation dynamics, suggesting in turn an at least partial induced-fit (active state induction) mechanism where adrenaline binding facilitates the receptor’s transition toward active-like conformations. This observation, if confirmed by further simulations and kinetic experiments, might contribute to the ongoing debate in the GPCR field regarding whether agonists exert their effects through *conformational selection* or *induced-fit mechanisms* [15].

### Modulation of ADRB1’s micro-switches

When we compare the active and inactive structures of a receptor, we see more localized changes in specific regions. These ‘micro-switch’ changes involve events like specific residue-to-residue contacts and side-chain rotations. Of the more than 90 state determinant micro-switches reported across different GPCR classes, only 4 seem to shared among all the classes [15]. Among the most critical micro-switches involved in ADRB1 activation, the PIF motif (i.e., I^3.40^, P^5.50^, and F^6.44^) plays a fundamental role in linking ligand binding to intracellular conformational changes (see Fig. 2a). In the apo-ADRB1 simulations, we observe a progressive reduction in the P^5.50^-F^6.44^ distance, shrinking from 1.3 nm in the inactive state to approximately 0.9 nm in the active state, marking a structural rearrangement that accompanies receptor activation. Conversely, in the holo-ADRB1 simulations, the P^5.50^-F^6.44^ distance oscillates stably around 0.9 nm throughout the whole Conformational Change CV, suggesting that the presence of adrenaline preconfigures the PIF motif into an activation-prone conformation. A similar trend is observed in the P^5.50^-I^3.40^ distance, which fluctuates between 0.4 and 0.7 nm in apo-ADRB1, reflecting the greater conformational flexibility of the receptor in the absence of the agonist. Instead, in the holo-ADRB1 simulations, this distance remains stably locked at 0.5 nm, even in conformations that correspond to the inactive state. These observations are consistent with an adrenaline facilitated shift of ADRB1’s conformational equilibrium toward an active-like state en route to a full activation.

**Figure 2:**
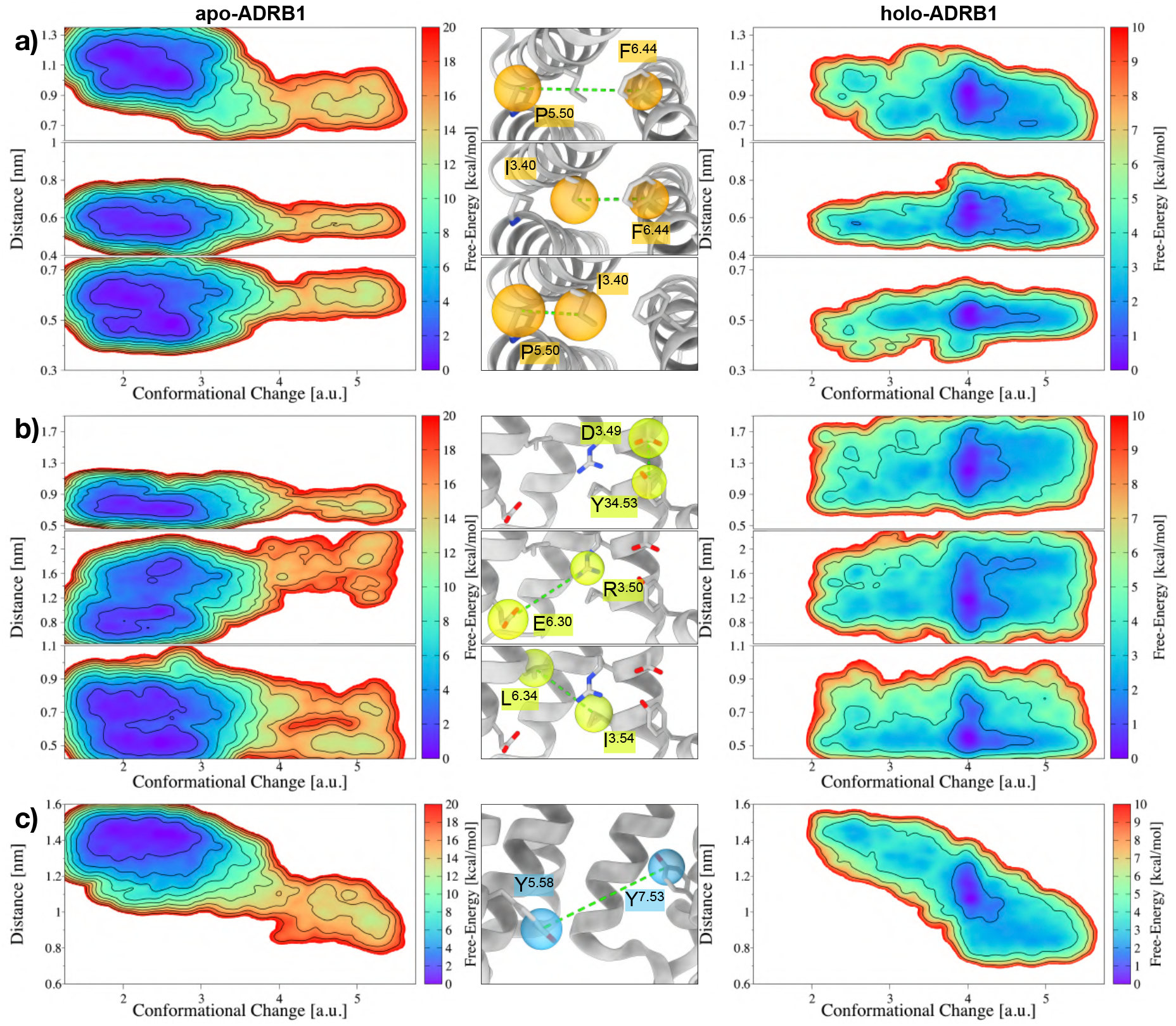
Analyses of ADRB1’s microswitches during its apo- and ligand-induced activation. **a)** 2D FES as a function of the Conformational Change CV and the respective Distance CVs for the residues of the PIF motif (P^5.50^-F^6.44^, I^3.40^-F^6.44^, and P^5.50^-I^3.40^). **b)** 2D FES as a function of the Conformational Change CV for the DRY motif (D^3.49^-Y^34.53^), the ionic lock (R^3.50^-E^6.30^), and an additional hydrophobic contact in the proximity of the DRY motif (I^3.54^-L^6.34^). **c)** 2D FES as a function of the Conformational Change CV for the YY motif (Y^5.58^-Y^7.53^). On the left side of (**a**), (**b**), and (**c**), we reported the average 2D FES of the 3 apo-ADRB1 OneOPES simulations, while on the right side the average 2D FES of the 3 holo-ADRB1 OneOPES simulations. Isolines are drawn every 2 kcal/mol.

In addition to the PIF motif, another set of key micro-switches that play a crucial role in ADRB1 activation is the DRY motif, consisting of D^3.49^, R^3.50^, and Y^3.51^ (see Fig. 2b). This conserved triad is involved with the surrounding amino acids in stabilizing the inactive state of GPCRs, and undergoes a significant rearrangement during receptor activation. Our analysis reveals that in the apo-ADRB1 simulations, the D^3.49^-Y^34.53^ hydrogen bond remains consistently engaged throughout the entire range of the Conformational Change CV. However, in the holo-ADRB1 simulations, the D^3.49^-Y^34.53^ distance becomes significantly more dynamic, fluctuating across a broad range of values between 0.9 nm and 1.7 nm. This increased mobility suggests that the presence of adrenaline weakens the constraints imposed by the DRY motif, allowing for a more flexible conformational landscape and facilitating the transition toward activation.

A similar trend is observed in the ionic lock, a crucial interaction between R^3.50^ and E^6.30^ that plays a well-established role in GPCR activation by stabilizing the inactive conformation. In the apo-ADRB1 simulations, the R^3.50^-E^6.30^ distance follows a well-defined activation pathway: in the inactive state, the salt bridge between R^3.50^ and E^6.30^ is broken, as displayed in the minimum at CC ∼2.5 and Distance ∼1.6 nm. This is a prerequisite for subsequent conformational rearrangements, allowing ADRB1 to transition toward its active state. However, in the holo-ADRB1 simulations, the R^3.50^-E^6.30^ interaction is far more mobile, displaying a broader distribution of values rather than following a single, well-defined reaction coordinate. Once again, the collected data suggest that the presence of adrenaline disrupts the tight structural constraints of the inactive state, increasing the receptor’s conformational plasticity and promoting alternative activation pathways.

Lastly, another crucial micro-switch that plays a fundamental role in ADRB1 activation is the YY motif, involving Y^5.58^ and Y^7.53^ on the intracellular side of the receptor. This motif is a well-known conserved structural element in class A GPCRs, often implicated in the stabilization of receptor conformations and in the allosteric regulation of intracellular signaling. As displayed in Fig. 2c, our analysis reveals that the evolution of the Y^5.58^-Y^7.53^ distance remains largely unperturbed by the presence of adrenaline, hinting that this specific interaction follows a similar activation pathway in both apo- and holo-ADRB1 simulations, aside from the shift of the deepest minimum in the 2D FES from CC ∼2.5 towards CC ∼4. The collected data may suggest that the YY motif does not exhibit significant structural alterations upon adrenaline binding. Nevertheless, it is important to stress that the YY motif is intrinsically linked to the hydration dynamics of ADRB1’s intracellular cavity. The Y^5.58^-Y^7.53^ interaction plays a pivotal role in controlling the influx and organization of water molecules, which, in turn, can significantly impact receptor activation by modulating key conformational transitions [51, 12]. In the pursuit of fully understanding the mechanistic effects of adrenaline binding, a comprehensive assessment of the hydration dynamics within ADRB1 has been carried out and is discussed in the subsequent sections of the manuscript.

### Hydration of ADRB1’s intracellular side

Changes in conserved water-mediated allosteric networks have been reported to play a crucial role in GPCR activation and signal transduction, including in ADRB1 [48, 51, 12, 49]. We analyzed the water occupancy and dynamics in both the apo- and adrenaline-bound states emerging from our OneOPES simulations, focusing on the intracellular side where conformational changes critical for G-protein coupling occur. To this end, we re-weighted the accumulated bias potentials on a 2D FES function of the Conformational Change CV and, in this case, the “Hydration” CV, defined by the position of a dummy atom located between residues Y^5.58^ and Y^7.53^ (for additional details, please refer to the “Methods” section and Fig. S1). This approach allowed us to quantify the relationship between hydration dynamics and ADRB1’s conformational transitions, providing valuable insights into the structural features of the conformational ensembles associated with each basin of the CC against the hydration FES.

Regarding the data collected on the apo-ADRB1 simulations, we observed the presence of a single well-defined free-energy minimum at CC ∼5 and a Hydration value in the ∼0.5-1.5 range, as reported in Fig. 3a. The analysis of the conformation pool corresponding to this basin let us identify transient 1-2 water molecules occupying the intracellular cavity. Such water molecules primarily interact with residues Y^5.58^ and Y^7.53^ of the YY-motif, forming short chains of hydrogen bonds between the two tyrosines mediated by water molecules. Conversely, the adrenaline-bound form exhibited a higher and more variable hydration pattern (e.g. up to 6 molecules), suggesting that ligand binding stabilizes a more hydrated environment (see Fig. 3b). Indeed, by extracting the conformations belonging to the lowest basin (i.e., CC ∼4 and Hydration ∼2.5), it was possible to observe a variety of water chains between Y^5.58^ and Y^7.53^, with the most frequent network being composed of 3 and 4 water molecules.

**Figure 3:**
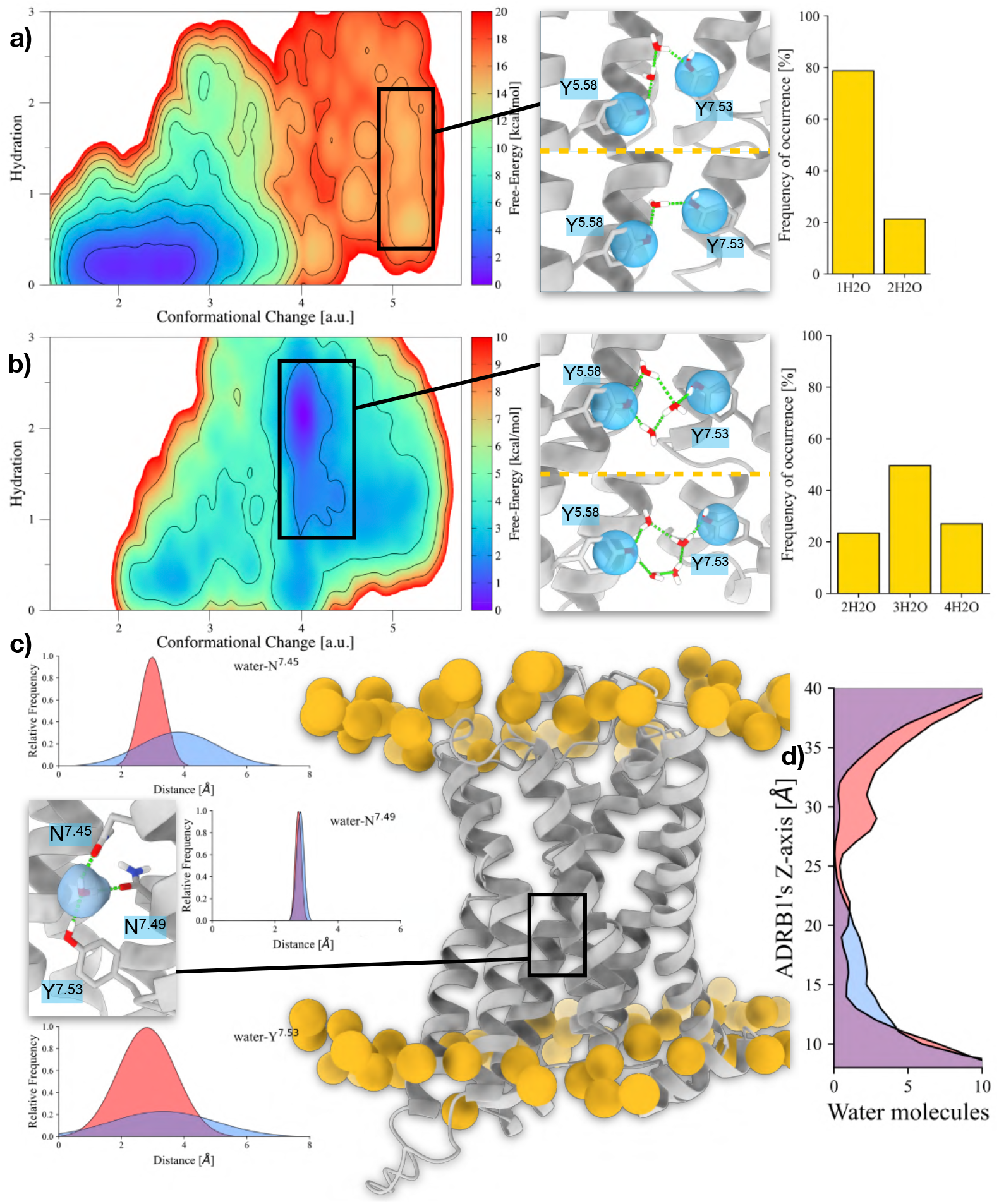
Differences in the internal hydration sites in apo- and holo-ADRB1. **a-b)** 2D FES associated with the hydration of (a) apo- and (b) holo-ADRB1’s YY-motif during the GPCR activation. On the right side, insets displaying the water molecules connecting Y^5.58^ and Y^7.53^ and the relative frequency of each hydration state in the selected basin. **c)** Average distances of N^7.45^ and N^7.53^ with respect to the closest water molecule to N^7.49^ in the apo- and holo-ADRB1 active basins. The image on the right depicts ADRB1 and the phosphorus atoms in the surrounding membrane. The inset displays the water molecule caged between N^7.45^, N^7.49^, and Y^7.53^. **d)** Distribution of water molecules along ADRB1’s z-axis. Values coming from apo- and holo-ADRB1’s active basins are colored in transparent red and blue, respectively.

To understand the underlying reason behind the differing hydration levels induced by adrenaline binding, we recognized that a purely numerical evaluation of the free-energy differences between the apo and holo conformations of ADRB1 would not be comprehensive. While the 2D FES provided critical insights into the hydration landscape, inspecting the structural impact of ligand binding in greater detail became imperative. We started with checking the overall cavity surrounding the YY-motif. This analysis let us observe the presence of structural water in the proximity of Y^7.53^. As depicted in Fig. 3c, the conformational ensemble extrapolated from apo-ADRB1’s active basin is characterized by a rigid network established by N^7.45^’s and N^7.49^’s carboxamides with Y^7.53^’s hydroxyl, which all insist on a single water molecule. In holo-ADRB1, these interactions are way looser, leading to an increased hydration of the intracellular side of ADRB1 (see Fig. 3d).

These analyses delivered a preliminary rationale for the differences in the intracellular activity between apo- and adrenaline-bound receptors, yet it does not clarify the consequences of ligand binding on ADRB1. So, we sought to reconstruct the cascade of structural rearrangements initiated by adrenaline at the extracellular side and how these propagate through the GPCR to influence the intracellular hydration environment. To achieve this goal, we compared the average pairwise distance of each residue pair in the apo- and holo-ADRB1 conformational pools. By mapping the shifts in residue pair distances, we could identify regions of the receptor undergoing significant rearrangements, which provided a mechanistic link between the extracellular binding event and its downstream effects on intracellular hydration.

As shown in Fig. 4, the binding of adrenaline in ADRB1’s orthosteric site perturbs the hydrophobic packing between F^6.52^ and W^6.48^, closely stacked in apo-ADRB1’s active state. At the same time, the positively charged ammine of adrenaline forms a salt bridge with D^3.32^, which in turn stabilizes the charge-enforced H-bonds with Y^7.43^. This first layer of interactions of the adrenaline propagates onto the same amino acid, i.e. N^7.45^. Indeed, in apo-ADRB1’s active state N^7.45^’s amide is stabilized by an H-bond with W^6.48^’s side-chain. Conversely, the displacement of W^6.48^ enhances the flexibility of N^7.45^ in holo-ADRB1, with the latter residue more prone to bind to G^7.42^’s carbonyl oxygen, in closer proximity due to Y^7.43^’s afore mentioned binding with D^3.32^.

**Figure 4:**
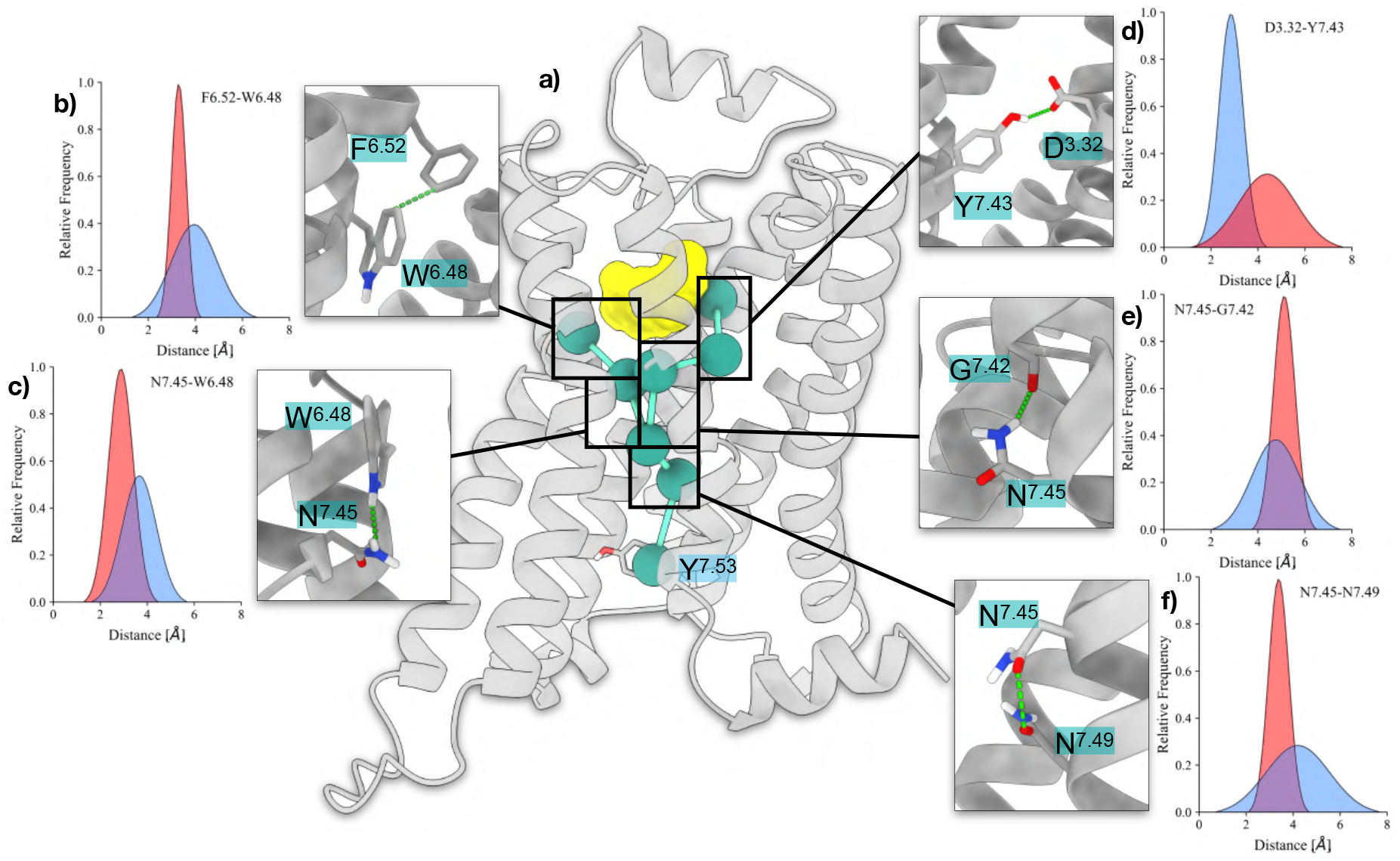
Schematic depiction of the effects of adrenaline binding at ADRB1 and the resulting propagation of structural changes through the receptor. **a)** Network of intraprotein contacts that initiates a series of conformational changes extending to the intracellular side upon adrenaline (yellow) binding. **b-f)** Average pairwise distances between the pair of residues F^6.52^-W^6.48^ (**b**), N^7.45^-W^6.48^ (**c**), D^3.32^-Y^7.43^ (**d**), N^7.45^-G^7.42^ (**e**), and N^7.45^-N^7.49^ (**f**). Values collected from apo- and holo-ADRB1’s active basins are colored in transparent red and blue, respectively.

Overall, this cascade of events led to an increase of the distance between N^7.45^ and N^7.49^, destabilizing the water-mediated interaction established between N^7.45^, N^7.49^, and Y^7.53^ presented in the previous paragraph, and favoring the entrance of additional water molecules in the intracellular side of ADRB1. These suggestions are further corroborated by the work of Chen et al. [49] that, during the preparation of the present manuscript, engineered de-novo highly signaling GPCRs by optimizing their intraprotein water network (centered on W^6.48^, N^7.45^, N^7.49^, and Y^7.53^), supporting our findings that the interaction of water molecules with polar residues can facilitate activation-prone movements in GPCRs.

### Allosteric modulation in the Sodium binding site

The sodium binding site of GPCRs is a conserved structural feature that plays a key role in regulating receptor activity and dynamics. In ADRB1, this site is located near the conserved D^2.50^ residue, which serves as a primary coordinating partner for sodium [52, 53]. Notably, D^2.50^ is also known to become protonated during receptor activation, a mechanism that stabilizes the active state and prevents sodium coordination. To investigate these two scenarios (namely, the effects of sodium binding and D^2.50^ protonation on ADRB1 activation) we carried out two novel sets of OneOPES simulations, each consisting of three independent replicas: apo-ADRB1-Na^+^, where we explicitly modeled sodium binding, and apo-ADRB1-ASPH, where D^2.50^ was protonated.

For apo-ADRB1-Na^+^, we extended our OneOPES sampling strategy to include a dedicated set of CVs that capture Na^+^ movement across the orthosteric cavity of the receptor (see Fig.5a and Tab. S2). Specifically, we introduced a *NaPATH* CV to track Na^+^’s motion from the extracellular side to the sodium binding site and a *NaWater* CV to monitor the hydration of sodium binding site itself (see Fig. S5). Reweighting the accumulated bias potential onto the Conformational Change CV and averaging the results over the three replicas, we observed that sodium binding markedly stabilizes the inactive state of ADRB1. Notably, we identified a significant energy barrier around milestone 5 of the NaPATH CV, where the ion must overcome a tightly packed hydrophobic cluster formed by M^2.53^, V^3.36^, and W^6.48^ (see Fig. 5b). This suggests that sodium entry is highly restricted at this point, reinforcing the notion that hydration dynamics and local structural rearrangements play an important role in sodium accessibility. Our free-energy calculations also indicate that ADRB1 activation is significantly disfavored in the presence of sodium, with a ΔG of activation of 15 ± 1 kcal/mol, further supporting experimental results (e.g. ^19^ *F* NMR, ^23^ *Na* NMR [53]) highlighting that sodium acts as an allosteric inverse agonist by stabilizing the receptor in its inactive conformation (see Fig. 5c and 5d).

**Figure 5:**
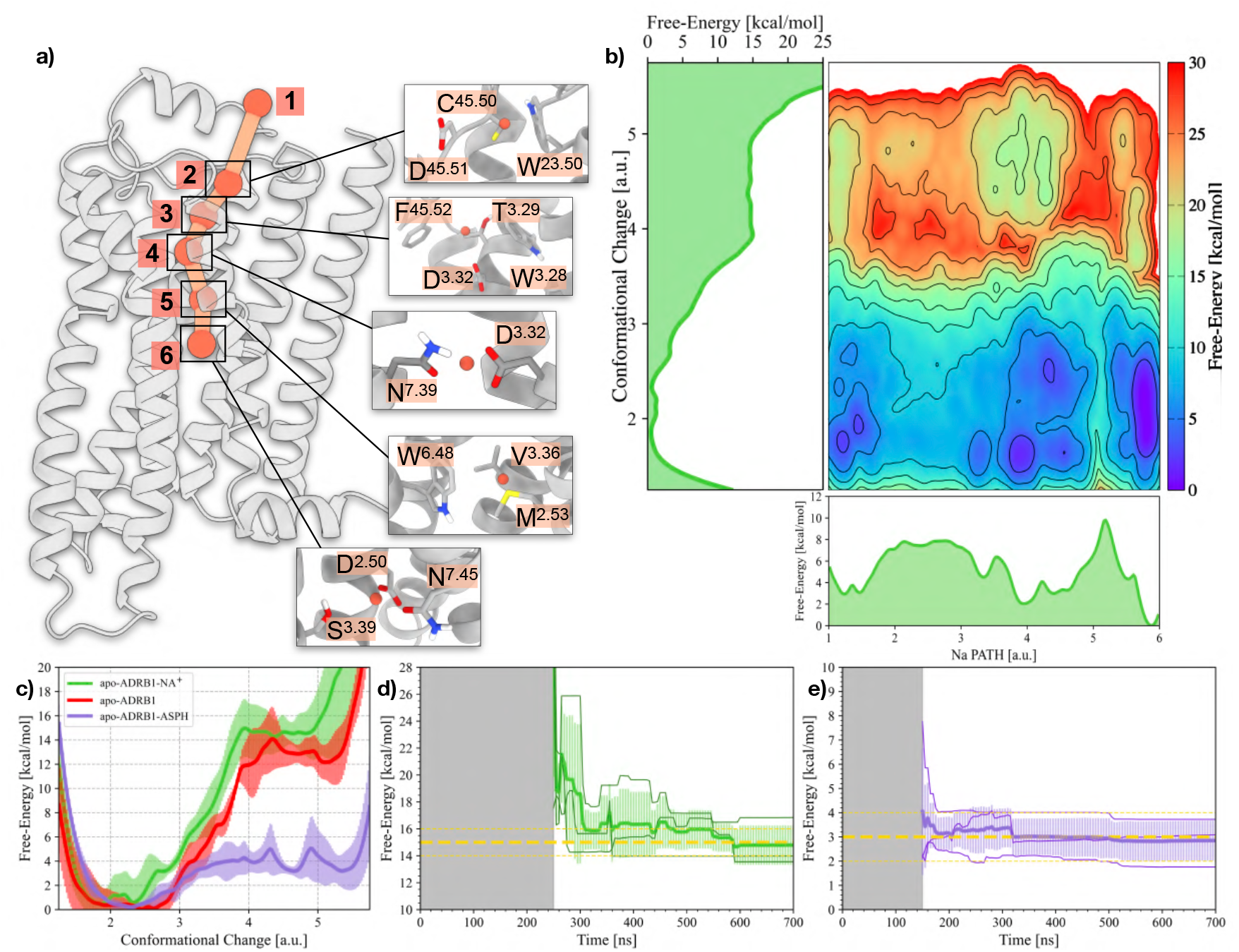
Allosteric modulation of the Na^+^ binding cavity. **a)** Schematic representation of the Na^+^ translocation pathway (NaPATH CV) within ADRB1. Milestones along the pathway are indicated as orange spheres and shown with inset enlargements for clarity. **b)** 2D FES illustrating the relationship between Na PATH (x-axis) and ADRB1’s Conformational Change CV (y-axis). 1D FES projections for the Na PATH and Conformational Change are also shown, providing insights into the energetic landscapes of these processes. **c)** 1D FES as a function of the Conformational Change CV for the apo-ADRB1, apo-ADRB1-NA^+^, and apo-ADRB1-ASPH systems. **d-e)** Free-energy difference between inactive and active states in the apo-ADRB1-NA^+^ (d) and apo-ADRB1-ASPH (e) systems over time. The average value is shown as a yellow dashed line. Error bars of 1 kcal/mol appear as yellow dashed lines.

Conversely, in the apo-ADRB1-ASPH simulations, where D^2.50^ was protonated, we applied the same OneOPES sampling strategy used for apo-ADRB1. The results revealed a significant population shift in the activation landscape: neutralization of D^2.50^ dramatically shifts the receptor toward the pre-active and active state, reducing the ΔG of activation to only 3 ± 1kcal/mol and making the pre-active and active states very close in energy (see Fig. 5c and 5e). This finding aligns with the known role of D^2.50^ protonation in receptor activation, further confirming that sodium binding and D^2.50^ protonation have opposite effects on the conformational equilibrium of ADRB1.

In physiological conditions, we can hypothesize that the D^2.50^ is initially protonated, facilitating the dynamical equilibrium with pre-active and even active states. In these states the Na^+^ has a lower barrier to enter in the conserved sodium pocket and, once there, it ‘locks’ the receptor in an inactive state until an agonist binds to the orthosteric site. In this regard, previous studies suggested that the presence of an agonist may then facilitate sodium egress towards the intracellular environment, potentially cutting down the free-energy cost associated with the full activation process [53]. For additional details regarding the apo-ADRB1-ASPH simulations, please refer to Fig. S6 and S7.

## Discussion

The ADRB1 receptor is an important pharmacological target playing a crucial role in cardiovascular regulation [54, 55, 56]. Our study aimed to leverage a novel, efficient and reproducible enhanced sampling technique, OneOPES, and to develop tailored collective variables to characterize the functional dynamics of GPCRs of great biomedical importance such as ADRB1. Through its combination of replicas exchange, thermal ramp, and multi-CVs, OneOPES has shown its ability to provide fully converged and reproducible results in a shorter simulation time compared to traditional methods, making it a powerful tool for studying GPCR activation. The tailored CVs developed here can be adapted and used to investigate the activation of other GPCRs. The reliability of our results is supported by the consistency observed across three independent multiple replica simulations for each different state (apo, holo, protonated), ensuring the reproducibility and robustness of our conclusions. Due to the high computational cost, the error bars on the calculated free energy profiles have in the past been approximated by methods such as block analysis. Here, for the first time for such complex systems, the computational efficiency of our approach allows us to obtain them directly from independent simulations. The computed free energy profiles for the apo and holo receptors are in line with high-resolution NMR data [51, 12] and provide details not accessible through other approaches.

In addition to providing key insights into the receptor’s activation pathway, particularly the role of water molecules in facilitating structural rearrangements, our study offers a comprehensive comparison of ADRB1 activation in both the apo and adrenaline-bound states. Our detailed analysis of the role of water in the activation mechanism is in line with previous reports [48, 49], and complements them with unprecedented details of the step-by-step role of water-mediated allosteric networks along the activation reaction coordinate. This comparative analysis of free-energy differences lays the groundwork for the rational design of novel agonists and antagonists. Future drug discovery efforts can leverage the framework provided by our simulations to assess the physiological activity of potential ligands computationally, significantly reducing the occurrence of false positives that would otherwise require costly experimental validation.

By streamlining the discovery process and improving the accuracy of ligand-receptor interaction predictions, OneOPES has the potential to accelerate the development of next-generation drugs targeting ADRB1. This method not only enhances our understanding of GPCR activation but also opens new avenues for designing more effective and selective therapies for cardiovascular diseases, ensuring better patient outcomes.

## Methods

### Systems preparation

The apo- and holo-structures of ADRB1 were generated starting from PDB ID: 7BVQ and 7BTS [36], respectively. The apo structure (PDB ID: 7BVQ) encompasses only a portion of the whole human ADRB1 sequence, notably residues S^1.28^ to S^5.74^ and V^6.26^ to C^8.59^. In the experimental structure, the ends of the sequences S^5.74^ and V^6.26^ were artificially linked, resulting in a shorter intracellular loop 3 (ICL3) compared to the native sequence. Regarding the holo-structure, the ICL3 is unresolved in PDB ID: 7BTS (from V^5.69^ to A^6.27^). For this reason, the missing patch was taken from 7BVQ and merged with the experimental structure. For both apo- and holo-ADRB1, the couples C^3.25^-C^45.50^ and C^45.43^-C^45.49^ were bound with a disulfide bridge. As described in previous studies [36], E^3.41^ was protonated because it directly contacts the phospholipid tails and predominantly exists in its neutral form.

The complexes thereby obtained were embedded into a simple phospholipid bilayer (POPC/CHL 80:20) using CHARMM-GUI [57], and solvated with the TIP4PD water model (salinity of 0.15 M NaCl). The N-terminus and C-terminus of both ADRB1 structures were capped with acetyl and metil-amino protecting groups, respectively. The DES-Amber force field was employed [58] in the MD engine GRO-MACS 2023 [59]. Each simulation box underwent a thermalization cycle with decreasing time-dependent restraints on heavy atoms, in order to relax unphysical bond lengths and retain the overall structure of the protein and the membrane components. All systems experienced the following protocol: 1 ns of NVT simulation followed by 1 ns of NPT simulation for each temperature, starting from 100 K until 300 K with steps of 50 K. The integration step was set to 2 fs. Coulomb and van der Waals interactions were cut-off at a distance of 1.0 nm. The particle-mesh-Ewald (PME) method was used to treat the long range electrostatic interactions [60]. The temperature was set at 300 K and controlled with the V-rescale thermostat [61], whereas the pressure was fixed at a reference value of 1 bar with the semi-isotropic C-rescale barostat [62].

### OneOPES MD simulations

A CV-based enhanced sampling method was adopted to investigate the conformational change of apo- and holo-ADRB1. Notably, we employed the OneOPES sampling scheme [39], a derivative technique of the “On-the-fly probability enhanced sampling” (OPES) algorithm in its “Explore” flavor [63]. In OneOPES, a replica-exchange framework of 8 independent trajectories is set up to ensure the exploration and the convergence of the FES under investigation. Such replicas are divided into two groups, i.e., a convergence-dedicated replica (replica **0**) and seven exploratory trajectories (replicas **1-7**). They all share OPES Explore as the main sampling engine carried out on a set of leading CVs. Replicas **1-7** are progressively heated (up to 335 K) thanks to OPES Expanded (OPES MultiT, hereafter), to ease overcoming hidden degrees of freedom [64]. To investigate the activation mechanism of ADRB1, we resorted to two tailored CVs. Among them, the main one is the “*PATH*” CV, a transition path composed of 13 protein conformations (i.e., milestones) representative of the conformational change between the inactive (milestone=1) and the active (milestone=13) ADRB1 structures and equally spaced in RMSD [65]. While this CV is sufficient to drive the overall motion of the C*α*, full activation of GPCRs is largely driven by the rearrangements of some selected amino acids’ side-chains that goes by the name of “*micro-switches*”. In the current scenario, we selected three sets of micro-switches to enhance the quality of the sampling:

- **PIF:** between the residues P^5.50^, I^3.40^, and F^6.44^, facilitating the movement of TM6;
- **DRY:** located at the cytoplasmic end of TM3, between D^3.49^, R^3.50^, and Y^3.51^;
- **NPxxY:** a sequence motif found in TM7, composed of N^7.49^, P^7.50^, and Y^7.53^.

In this study, the distances between the elements of these motifs have been treated as individual CVs to effectively capture and encourage transitions between the active and inactive states of ADRB1.

Recent literature has highlighted the critical role of water molecules in the full activation of the ADRB1 GPCR, particularly their entry into the cytoplasmic side of the receptor. Notably, the presence of water molecules is necessary to bridge the interaction between Y^7.53^ of the NPxxY motif and Y^5.58^, facilitating the receptor’s activation. To support this mechanism, we included a “*Hydration*” CV between these two side chains in our study, aiming to favor the activation process by ensuring proper water-mediated interactions [66]. All of these auxiliary CVs have been implemented in the exploration-dedicated replicas.

For a complete description of the atoms involved in the above-mentioned CVs, please refer to Fig. S1. In this framework, the exchange frequency among the replicas was 5000 integration steps. Replicas **0-7** contain a layer of OPES Explore with a bias of 100.0 kJ/mol and a PACE (i.e., frequency of bias deposition) of 50000 steps. Moreover, replicas **1-7** underwent additional OPES Explore layers (OPES MultiCV) with a bias of 3.0 kJ/mol on the auxiliary CVs and a PACE of 100000 steps. We selected the following temperature range, i.e. [300K: 301K] for replica **1**, [300K: 303K] for replica **2**, [300K:306K] for replica **3**, [300K: 310K] for replica **4**, [300K: 317K] for replica **5**, [300K: 325K] for replica **6**, and [300K: 335K] for replica **7**. For clarity, a full description of the different CVs and parameters used for the OneOPES simulations has been reported on Tab. S1. To run the simulations the MD engine GROMACS 2023 patched with PLUMED 2.9 was employed [67]. Regarding the thermostat and the barostat, we used the same protocol described in the section “MD simulations”.

### Cluster analysis

Cluster analyses on the MD trajectories were performed using GROMACS’s *gmx cluster* routine, using the *gromos* algorithm. An RMSD threshold value of 2.0 Å was selected considering the number of generated cluster families and the similarity of protein conformations within a cluster family.

### Binding interface evaluation

To assess the interactions between the adrenaline and ADRB1 in the holo-ADRB1 MD simulations, we used the PLOT NA routine of the “Drug Discovery Tool” (DDT) to estimate the frequency of occurrence of contacts [68]. To analyze the binding interfaces, we followed the approach of previous studies and set a neighboring cutoff value of 3.5 Å between the ligand and the interacting residues.

## Data Availability

The input files to replicate all the simulations can be found on https://github.com/valeriorizzi/ADRB1_OneOPES.

## Acknowledgements

The work was financially supported by the Swiss National Science Foundation and Bridge funding schemes (project numbers: 200021 204795, CRSII5 216587, and 40B2-0 203628). We acknowledge the Swiss National Supercomputing Centre for supercomputer time allocations on Piz Daint (project ID: s1274). We wish to thank Ioannis Galdadas, Dorothea Gobbo and Maria Bzowka for useful discussions, especially on effective CV combinations.

