## Supplementary Material for "Revealing Water-Mediated Activation Mechanisms in the Beta 1-Adrenergic Receptor via OneOPES-Enhanced Free Energy Landscapes"

The data hereby reported are divided into the following sections:

- Supplementary Data 1: Details on the employed CVs and how they have been developed;
- Supplementary Data 2: Details on apo-ADRB1's activation PATH OneOPES simulations;
- Supplementary Data 3: Details on holo-ADRB1's activation PATH OneOPES simulations;
- Supplementary Data 4: Adrenaline's binding mode in the intermediate states;
- Supplementary Data 5: Details on the CVs designed to monitor $Na^+$;
- Supplementary Data 6: Details on the apo-ADRB1-$NA^+$ OneOPES simulations;
- Supplementary Data 7: Details on the apo-ADRB1-ASPH's OneOPES simulations.

### Supplementary Data 1

To investigate the conformational dynamics of the apo-ADRB1 and holo-ADRB1 systems, we employed a PATH CV (*p1.s* in Tab. S1 and Tab. S2) and a set of carefully designed CVs that capture key structural and hydration-related changes during receptor activation. These CVs were selected and constructed to comprehensively represent the molecular processes governing receptor function, including water-mediated transitions and residue rearrangements. In particular, the CVs were built following a coarse-grained-inspired approach, treating the entire side-chain of specific amino acids as putative dummy atoms to simplify and focus the representation on functionally relevant dynamics. For detailed atomistic descriptions, please refer to Fig. S1. To see how the different CVs are distributed across the OneOPES replicas, please refer to Tab. S1.

| Replicas | 0 | 1 | 2 | 3 | 4 | 5 | 6 | 7 |
|---|---|---|---|---|---|---|---|---|
| OPES Explore | p1.s | p1.s | p1.s | p1.s | p1.s | p1.s | p1.s | p1.s |
| OPES MultiCV 1 | - | **D1**,yywo | **D1**,yywo | **D1**,yywo | **D1**,yywo | **D1**,yywo | **D1**,yywo | **D1**,yywo |
| OPES MultiCV 2 | - | - | **D2**,yywo | **D2**,yywo | **D2**,yywo | **D2**,yywo | **D2**,yywo | **D2**,yywo |
| OPES MultiCV 3 | - | - | - | **D3**,yywo | **D3**,yywo | **D3**,yywo | **D3**,yywo | **D3**,yywo |
| OPES MultiCV 4 | - | - | - | - | **D4**,yywo | **D4**,yywo | **D4**,yywo | **D4**,yywo |
| OPES MultiCV 5 | - | - | - | - | - | **D5**,yywo | **D5**,yywo | **D5**,yywo |
| OPES MultiCV 6 | - | - | - | - | - | - | **D6**,yywo | **D6**,yywo |
| OPES MultiCV 7 | - | - | - | - | - | - | - | **D7**,yywo |
| OPES MultiT | - | 301K | 303K | 306K | 310K | 317K | 325K | 335K |

Table S1: Table depicting the different CVs and parameters used for the OneOPES simulations. On the rows "OPES Explore" and "OPES MultiCV" we report how the CVs have been arranged along the replicas. On the row "OPES MultiT", we display the highest temperature achieved along the trajectory starting from the temperature of the thermostat (i.e., 300 K).

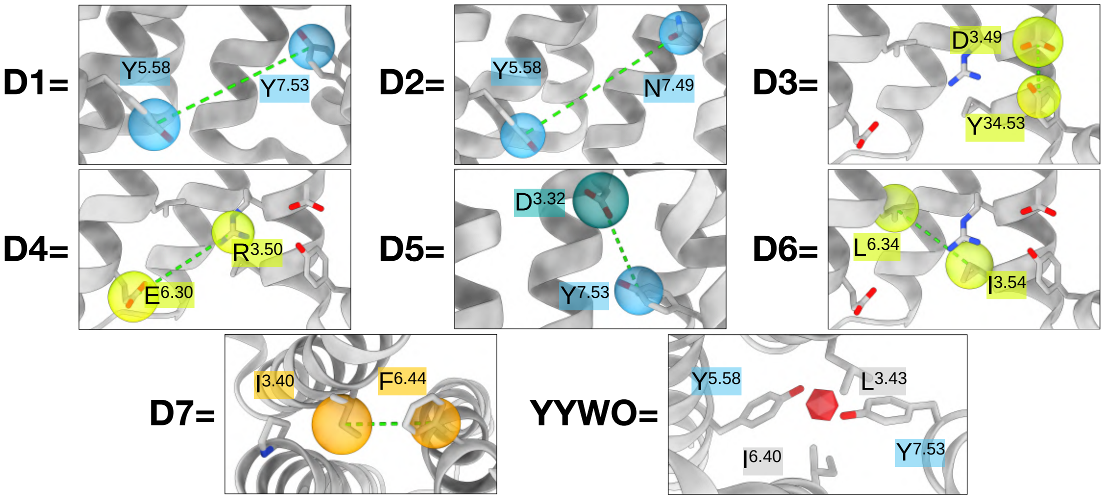


Figure S1: **Atomistic details of the CVs employed in the apo- and holo- ADRB1 OneOPES simulations.** For **D1-D7**, each CV is built as a distance between dummy atoms centered on side-chains. Each dummy atom is colored according to Fig. 3. For **YYWO**, the dummy atom built between L$^{3.43}$, I$^{6.40}$, and Y$^{7.53}$ and coordinating water molecules is represented as a red icosahedron.

#### Supplementary Data 2

The following section presents additional data collected from the three apo-ADRB1 OneOPES simulations to support our investigation. Specifically, we display the free energy as a function of the ADRB1 activation path, providing insights into the energy landscape of the receptor's conformational changes (see Fig. S2a). Additionally, we include three plots showing the sampling of the CVs "PATH", "Conformational Change", and "Hydration" across replica 0 of one apo-ADRB1 OneOPES simulation, highlighting the efficiency of the enhanced sampling and the distribution of key variables along the activation pathway.

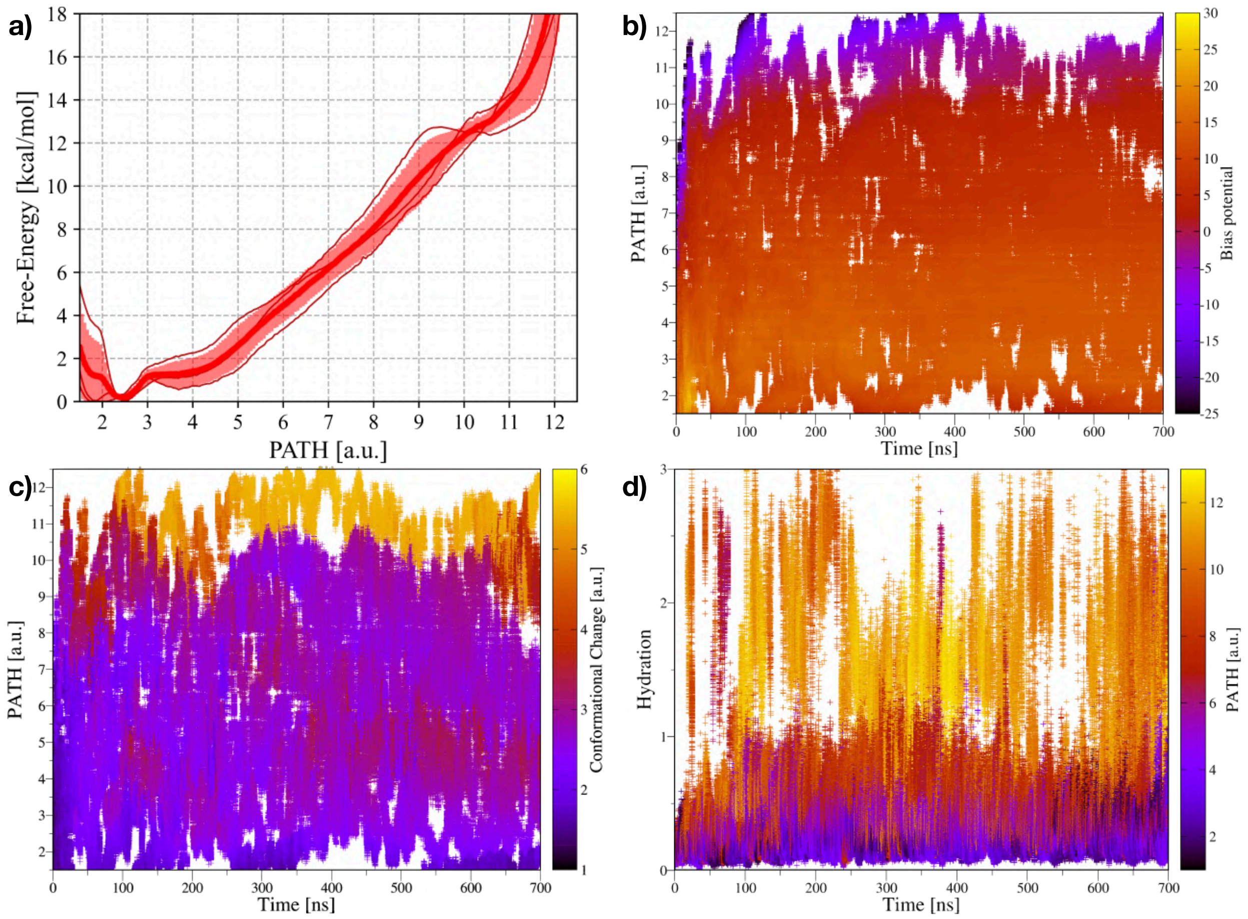


Figure S2: **Free-energy landscape and CV sampling in apo-ADRB1 OneOPES simulations.** **a)** Free-energy profile as a function of the ADRB1 activation PATH, averaged over three independent OneOPES simulations. The solid red line represents the mean free-energy, while the transparent red shading indicates the standard deviation. **b-c)** Sampling of the PATH CV as a function of time in replica **0**, with data points colored according to the accumulated bias potential and Conformational Change CV (panels **b** and **c**, respectively). **d)** Sampling of the Hydration CV as a function of time in replica **0**, with data points colored according to the PATH CV.

#### Supplementary Data 3

In the following section, we present additional data from the holo-ADRB1 OneOPES simulations to support our investigation. Notably, we display the free energy as a function of the ADRB1 activation path, providing insight into the GPCR in its ligand-bound state. Additionally, we include three plots showing the sampling of the CVs "PATH," "Conformational Change," and "Hydration" across replica **0** of one holo-ADRB1 OneOPES simulation.

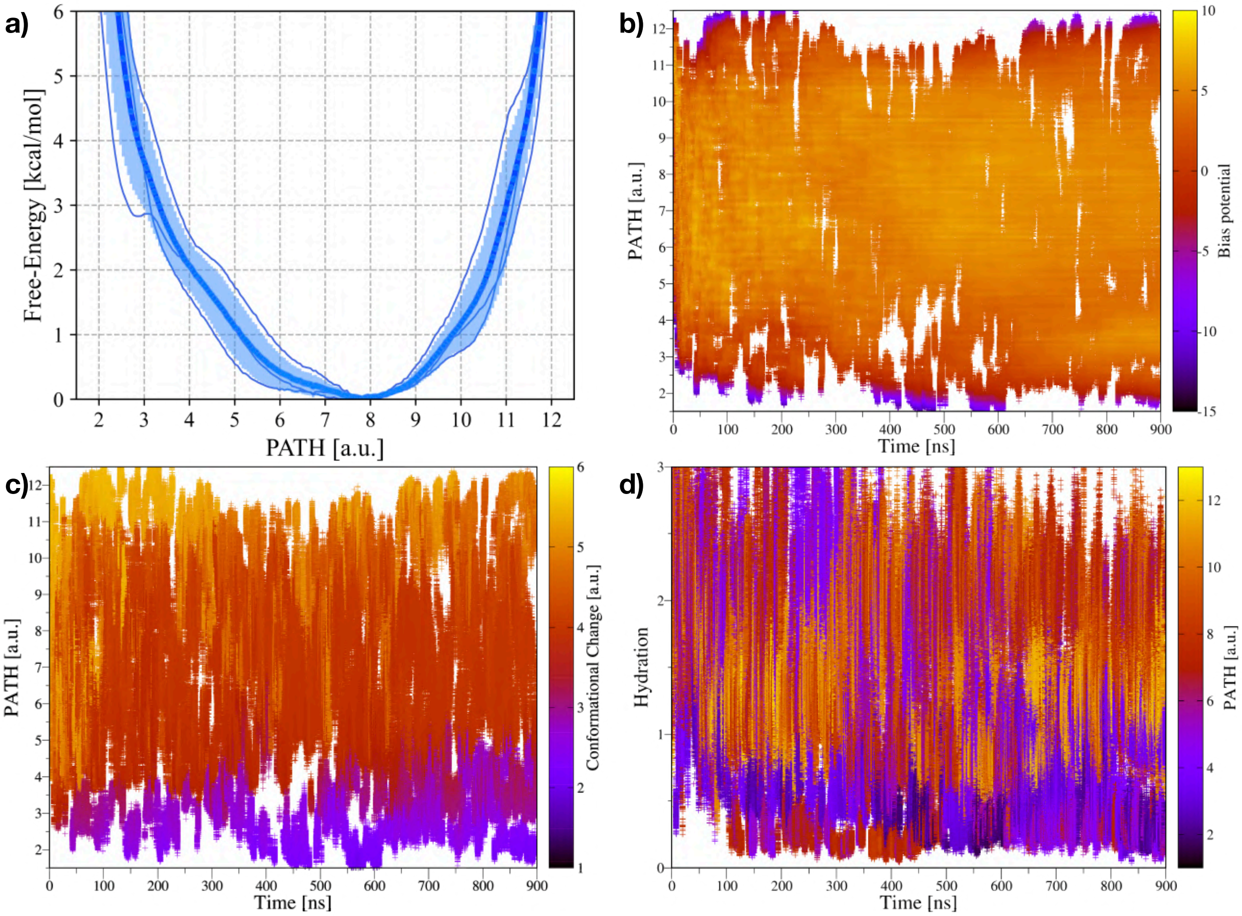


Figure S3: **Free-energy landscape and CV sampling in holo-ADRB1 OneOPES simulations. a)** Free-energy profile as a function of the ADRB1 activation path, averaged over three independent OneOPES simulations. The solid blue line represents the mean free-energy, while the transparent blue shading indicates the standard deviation. **b-c)** Sampling of the PATH CV as a function of time in replica **0**, with data points colored according to the accumulated bias potential and Conformational Change CV (panels **b** and **c**, respectively). **d)** Sampling of the Hydration CV as a function of time in replica **0**, with data points colored according to the PATH CV.

#### Supplementary Data 4

In Fig. S4, we present further insights into the conformational landscape and ligand-binding properties of ADRB1 in both apo and holo states. The first two panels illustrate the construction of the "*Conformational Change*" (CC) path, mapped onto the 2D FES built as a function of the RMSD relative to the inactive and active ADRB1 structures. The third panel compares the adrenaline binding mode observed in the pre-active state, identified from holo-ADRB1 OneOPES simulations, with the binding mode found in the crystallographic active structure (PDB ID 7BTS). Finally, the fourth panel presents a histogram reporting the frequency of occurrence of ADRB1 residues involved in adrenaline binding within the ensemble of pre-active state conformations, highlighting key interactions that stabilize this intermediate [1].

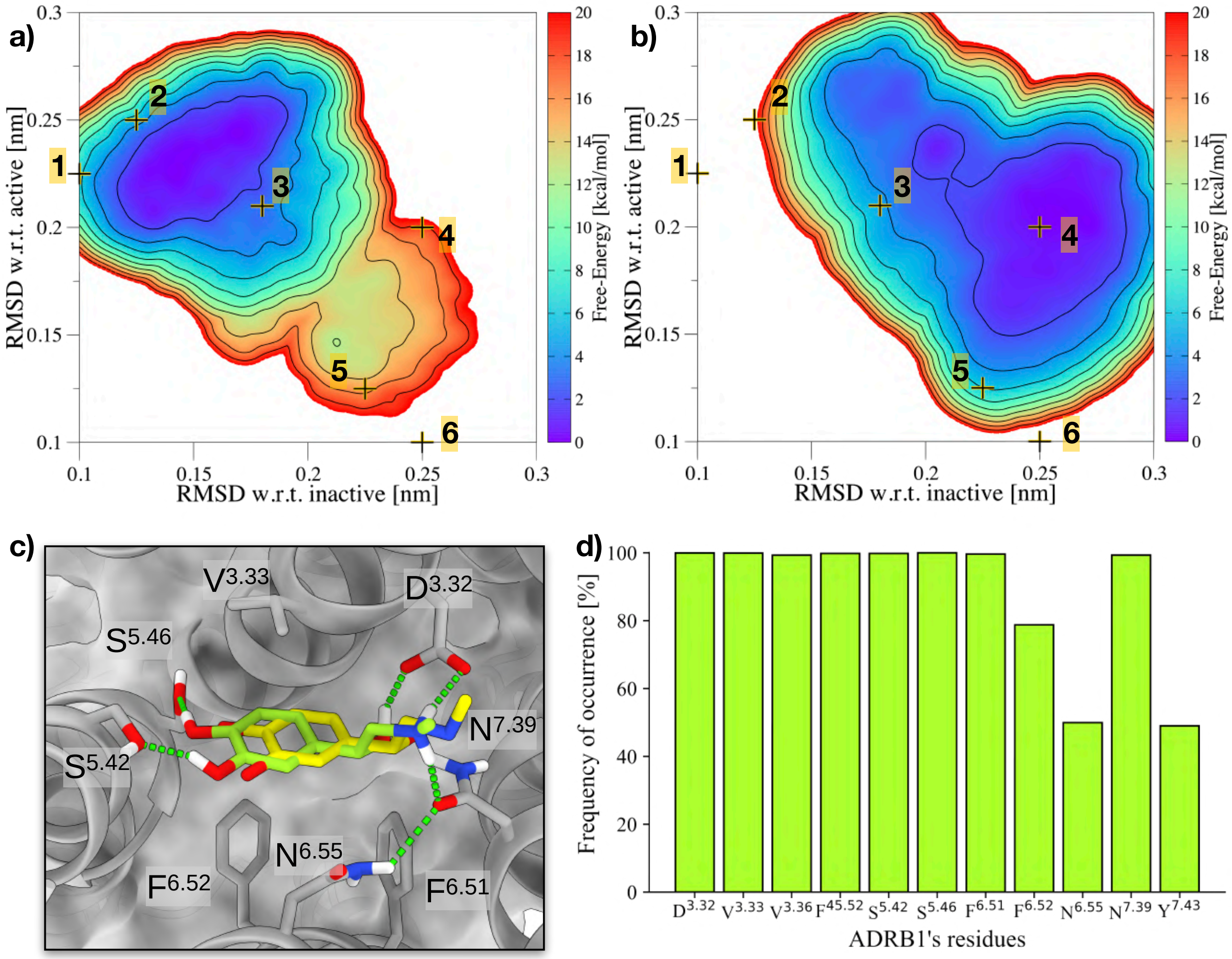


Figure S4: **Additional analyses of apo- and holo-ADRB1 OneOPES simulations. a-b)** Construction of the "Conformational Change" (CC) path, mapped onto the 2D FES built as a function of the RMSD relative to the inactive and active ADRB1 structures. **c)** Comparison of the adrenaline binding mode in the pre-active state (CC=4) with the binding mode observed in the crystallographic structure 7BTS. **d)** Histogram reporting the frequency of occurrence of ADRB1 residues involved in adrenaline binding within the ensemble of pre-active state conformations.

#### Supplementary Data 5

The following section reports additional technical details about the apo-ADRB1's OneOPES simulations in which we sample the motion of $Na^+$ ions towards the $Na^+$ binding cavity (i.e., apo-ADRB1-$NA^+$). To fully sample both the GPCR activation path and the $Na^+$'s translocation, the *NaWater* CV was designed to ease the hydration dynamics associated with specific regions of ADRB1's orthosteric binding site, as illustrated in Fig. 5a. This CV is expressed as a linear combination of water coordination numbers around three dummy atoms (denoted as **4**, **5**, and **6** in Fig. 6). In this way, *NaWater* allows it to capture the displacement of the hydration equilibrium between these regions, thereby serving as a meaningful descriptor for the ADRB1's conformational changes. The *NaWater* CV is defined as:

$$NaWater = (W_6 - W_5) - (W_4 - W_5)$$

where $W_i$ represents the number of water molecules coordinated to the dummy atom $i$. Coordination numbers were computed as in refs. [2, 3], with the difference that $Na^+$ ions were added to the group of water molecules. As illustrated in Fig. S5, this construction ensures the following behavior of *NaWater*:

- When the dummy atoms **4** and **6** are fully hydrated, *NaWater* assumes a value of +2;
- When the dummy atom **5** is fully hydrated, *NaWater* assumes a value of -2.

The intermediate values of *NaWater* correspond to varying degrees of hydration asymmetry between the regions defined by these dummy atoms. This behavior allows *NaWater* to serve as an intuitive descriptor for transitions associated with water-mediated conformational changes.

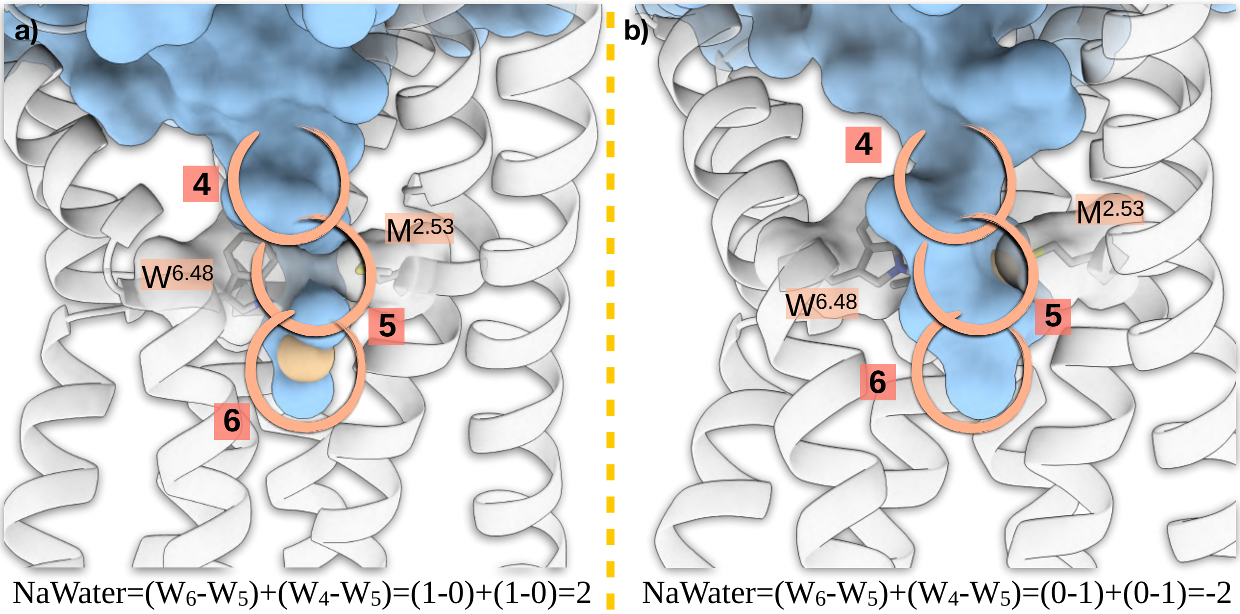


Figure S5: **Schematic descriptions of the *NaWater* CV. a)** Structural representation showing the hydrophobic residues $W^{6.48}$, $M^{2.53}$, and $V^{3.36}$, which form a barrier separating the water chambers around the NaPATH milestones **4** and **6**. This segregation maintains the sodium ion in the $NA^+$ binding site. **b)** Mechanism illustrating how water pumping across the hydrophobic residues $W^{6.48}$, $M^{2.53}$, and $V^{3.36}$ facilitates the transition of the sodium ion through the NaPATH milestone **5**. ADRB1 is represented in cartoon and colored in gray, whereas the water molecules are displayed as a continuous azure surface. $NA^+$ ion is represented as a yellow rigid sphere.

#### Supplementary Data 6

To investigate how the motion of $Na^+$ ions toward the $Na^+$ binding site affects the conformational dynamics of the apo-ADRB1 system, we performed novel OneOPES simulations using an updated set of carefully designed CVs that capture both the receptor's activation pathway and the motion of $Na^+$ ions. Specifically, the $Na^+$ ion motion was encoded within a newly developed path variable, *NaPATH*, which was designed to describe the ion's displacement along relevant interaction sites:

$$\text{NaPATH} = \frac{\Sigma_{i=1}^{6} i \cdot \exp[-\lambda(DNaPoint_i)^2]}{\Sigma_{i=1}^{6} \exp[-\lambda(DNaPoint_i)^2]}$$

where $DNaPoint_i$ is the distance of $Na^+$ from the NaPATH milestones represented in Fig. 6a. This variable was combined with the pre-existing CVs **D1–D7** and **YYWO**, as well as the previously introduced *NaWater* CV, which accounts for hydration changes critical to receptor activation. This combination of CVs allowed us to efficiently explore the receptor's free-energy landscape and identify key intermediates along the activation process. The distribution of CVs across the OneOPES replicas is reported in Tab. S2.

| Replicas | 0 | 1 | 2 | 3 | 4 |
|---|---|---|---|---|---|
| OPES Explore | p1.s,NaPATH | p1.s,NaPATH | p1.s,NaPATH | p1.s,NaPATH | p1.s,NaPATH |
| OPES MultiCV 1 | - | NaWater,yywo | NaWater,yywo | NaWater,yywo | NaWater,yywo |
| OPES MultiCV 2 | - | - | **D1**,yywo | **D1**,yywo | **D1**,yywo |
| OPES MultiCV 3 | - | - | - | **D2**,yywo | **D2**,yywo |
| OPES MultiCV 4 | - | - | - | - | **D3**,yywo |
| OPES MultiCV 5 | - | - | - | - | - |
| OPES MultiCV 6 | - | - | - | - | - |
| OPES MultiCV 7 | - | - | - | - | - |
| OPES MultiCV 8 | - | - | - | - | - |
| OPES MultiT | - | 301K | 303K | 306K | 310K |

| Replicas | 5 | 6 | 7 |
|---|---|---|---|
| OPES Explore | p1.s,NaPATH | p1.s,NaPATH | p1.s,NaPATH |
| OPES MultiCV 1 | NaWater,yywo | NaWater,yywo | NaWater,yywo |
| OPES MultiCV 2 | **D1**,yywo | **D1**,yywo | **D1**,yywo |
| OPES MultiCV 3 | **D2**,yywo | **D2**,yywo | **D2**,yywo |
| OPES MultiCV 4 | **D3**,yywo | **D3**,yywo | **D3**,yywo |
| OPES MultiCV 5 | **D4**,yywo | **D4**,yywo | **D4**,yywo |
| OPES MultiCV 6 | - | **D5**,yywo | **D5**,yywo |
| OPES MultiCV 7 | - | - | **D6**,yywo |
| OPES MultiCV 8 | - | - | **D7**,yywo |
| OPES MultiT | 317K | 325K | 335K |

Table S2: Table depicting the different CVs and parameters used for the apo-ADRB1-$NA^+$ OneOPES simulations. On the rows "OPES Explore" and "OPES MultiCV" we report how the CVs have been arranged along the replicas. On the row "OPES MultiT", we display the highest temperature achieved along the trajectory starting from the temperature of the thermostat (i.e., 300 K).

#### Supplementary Data 7

In this section, we present additional data collected from the three apo-ADRB1 OneOPES simulations in which the conserved $D^{2.50}$ residue was protonated (apo-ADRB1-ASPH hereafter). These simulations provide further insights into the receptor's conformational dynamics under altered protonation states. Fig. S6 presents macroscopic analyses of the apo-ADRB1-ASPH system, highlighting key structural and energetic features. Fig. S7 focuses on the behavior of critical micro-switches as a function of the Conformational Change CV, offering a detailed view of their role in receptor activation.

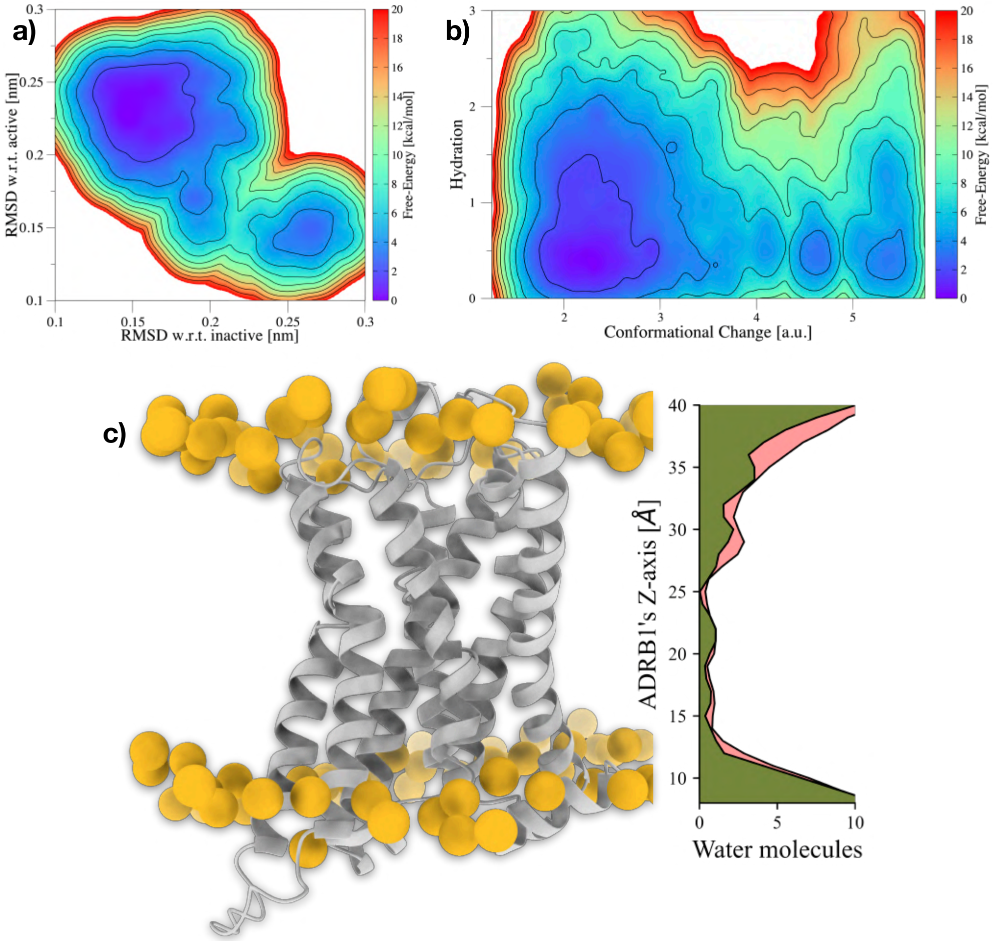


Figure S6: **Macroscopic analyses of apo-ADRB1 OneOPES simulations with protonated $D^{2.50}$.** **a)** 2D FES as a function of the RMSD with respect to the inactive and active conformations of ADRB1, illustrating the system's conformational landscape. **b)** 2D FES as a function of the Conformational Change and Hydration CVs, highlighting the relationship between receptor progression along the activation pathway and hydration dynamics. **c)** Distribution of water molecules along the z-axis of ADRB1, providing insights into hydration patterns within the receptor's transmembrane region. The profile collected from apo-ADRB1 is colored red, while the profile gathered from apo-ADRB1-ASPH is colored in green.

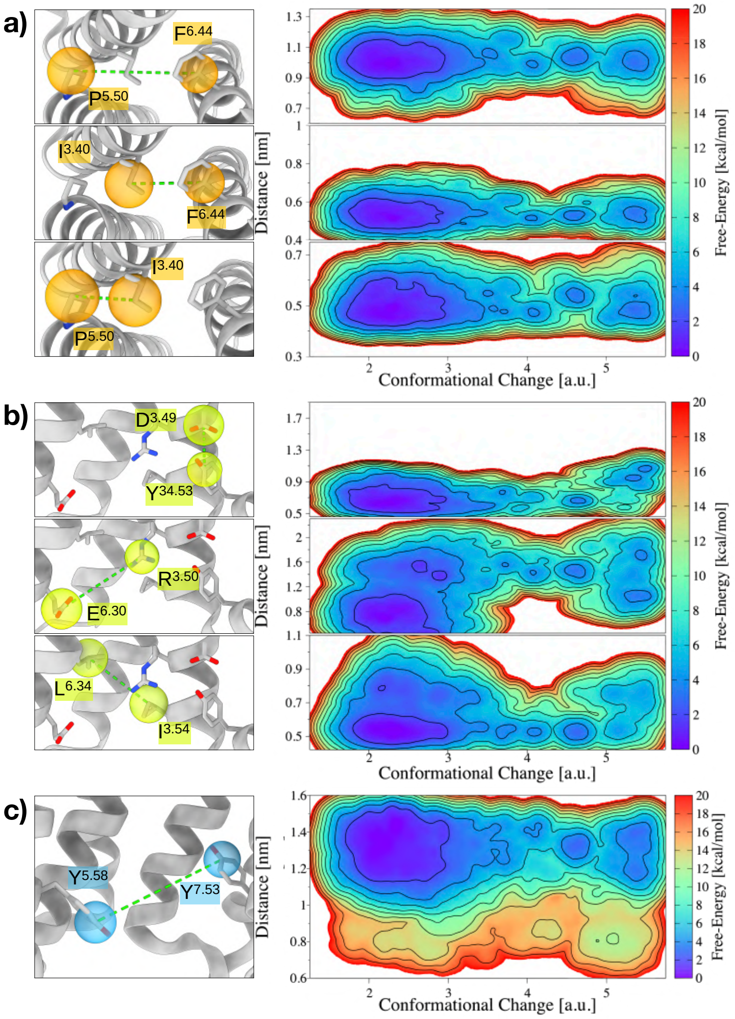


Figure S7: **Micro-switch analyses in apo-ADRB1-ASPH OneOPES simulations**. **a)** 2D FES as a function of the Conformational Change CV and the PIF distances, illustrating the coupling between receptor progression and the PIF micro-switch. **b)** 2D FES as a function of the Conformational Change CV and the DRY distances, highlighting the structural transitions associated with the DRY micro-switch. **c)** 2D FES as a function of the Conformational Change CV and the YY distance, providing insights into the role of the YY micro-switch along the activation pathway.
